# Improving face age prediction by using multiple-angle photos

**DOI:** 10.1101/2025.08.11.669799

**Authors:** Botond Bárdos-Deák, Csaba Kerepesi

## Abstract

Face photo-based age provides a cost-effective and readily accessible tool for biological age studies. However, face photo age models were usually trained on a single front-view photo per subject. Here, we hypothesized that face photo-based age prediction performance might be improved by using multiple photos of the same subject at the same time, captured from different angles. To test this hypothesis, we used an available dataset containing mugshots and developed age prediction models trained on (i) only front-view images, (ii) only the side-view images, and (iii) both front and side images. We found that accurate age prediction is possible using side photos despite the smaller facial area compared to front-facing photos (MAE = 3.1 years for the front-view and MAE = 3.7 years for the side-view images). The age prediction performance further improved by using two images from one person at the same time, captured from two different angles, front and side (MAE = 2.9 years). We found that subjects who age faster based on front-view face photos generally age faster based on side-view face photos. We also found that side-view models handle the rotation of the face better compared to the front-view model. In summary, we showed that two photos at different angles can improve age prediction and may provide a better approach to determining biological age for personalized medicine and rejuvenation studies.

## Introduction

Aging is a major problem of the 21st century^1^. Solving aging is a big scientific challenge where bioinformatics and artificial intelligence can contribute by developing age prediction models (i.e, aging clocks) that provide accurate measurement of aging and biological age^2–4^. In the last decade, different types of aging clock models were developed based on microscopic and macroscopic data for measuring the biological age of an individual. Aging clocks not only predicted age accurately but also better predicted all-cause mortality better than chronological age^5–10^. Hence, aging clocks suggest that some people age faster or slower than others, and accordingly, their biological age can be higher or lower than their chronological age. Age prediction models based on standard 2D face photos^11–13^ and 3D face imaging^14–16^ were also developed. In the current study, we focus only on the simpler and standard 2D face photo-based aging clock models that provide a low-cost, readily accessible biological age approach. Recently, we showed that face photo-based age acceleration calculated by artificial intelligence is predictive of all-cause mortality^17^. Others showed that face photo-based age predicted mortality of cancer patients better than chronological age^18^. These results suggest that face photo-based age may measure biological age and provides a cost-effective and readily accessible tool for personalized medicine and rejuvenation studies. However, face photo-based aging clocks typically were trained on a single photo per person, usually captured from the front view. Here we investigate whether face photo-based age prediction performance can be improved by using two photos of the same person at the same time, captured from different angles, one from front-view and one from side-view. We hypothesize that two photos at different angles can improve age prediction and may provide a better approach for determining biological age.

## Results

### Improved face age prediction using both front-view and side-view facial photos

We tested whether face photo-based age prediction performance can be improved by using two photos of the same person at the same time captured from different angles (Fig. 1a). For this purpose, we collected available data of mugshot pairs (front-view and side-view face photos) of 54,295 Illinois state prisoners (the “Prisoner dataset” hereafter, see the Methods). We split the high-quality mugshots of the Prisoner dataset into training, validation, and testing sets by the subjects. Then we developed three separate models trained on (i) only front-view facial mug shots (*Front model*), (ii) only side-view facial mug shots (*Side model)*, and (iii) both front-view and side-view facial mug shots (*Front + Side model*). We developed another model by combining the predictions of the front model and side models (*Combined model)*. We trained the models on the training set, optimized using the validation set, tested the final models on the testing set predicting the age of the subjects. The *Front model* was more accurate than the *Side model* (MAE = 3.1 vs 3.7 years). However, using two images, the age prediction performance further improved (MAE = 3.01 years by the *Front+Side* model and MAE = 2.88 years by the *Combined model*, Fig. 1bc, Supplementary Table S1).

**Fig. 1.**
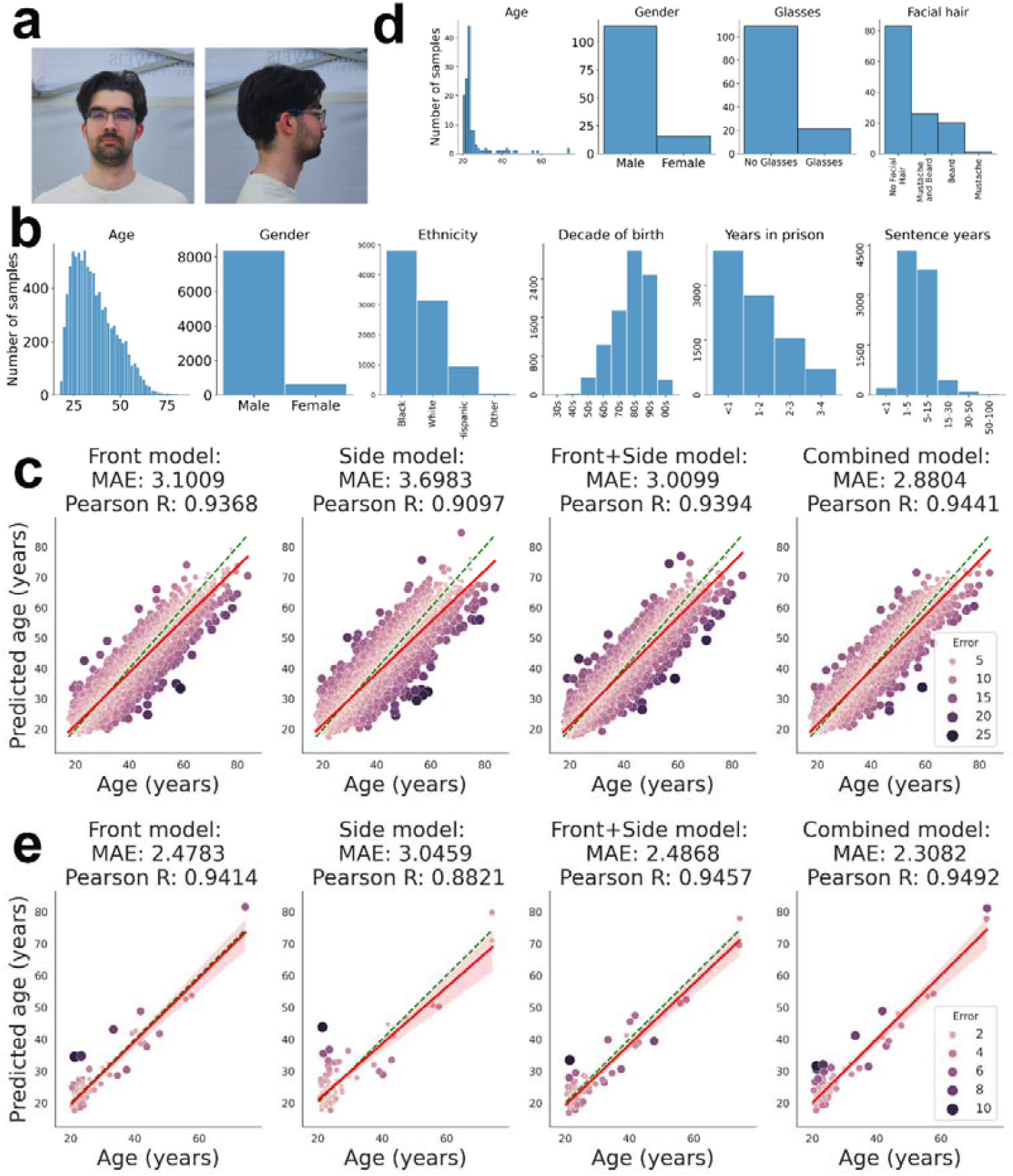
Improved face age prediction using both the front-view and side-view facial photos. **a**, An illustrative mugshot from the front (left) and side (right) of the first author of this study (the training and testing sets did not include these two pictures). **b**, Distribution of the attributes of the testing set of the Prisoner dataset. **c**, Age prediction performances of the face photo age models using the testing set of the Prisoner dataset. **d**, Distribution of the SCface dataset attributes. **e**, Prediction performances of the face photo age models on the external dataset, SCface.

We also tested the age prediction models on an external dataset, SCface^19^ containing high-quality photos of 130 subjects taken from different angles (Fig. 1d). We applied the *Front model* on the front-facing photos, the *Side model* on the side photos, as well as, the *Front+Slide* and *Combined models* on both photos (Fig. 1e, Supplementary Table S1). Again, the *Front model* was more accurate than the *Side model* (MAE = 2.48 vs 3.05 years), and the *Combined model* was the best (MAE = 2.31 years). Furthermore, we tested the *Front model* on two other independent external datasets, the IMDB-Clean and MORPH-2 (Fig. 2ab). The model generalized well in the MORPH-2, containing mugshots, however, did not perform well in the “in-the-wild” dataset of the IMDB-Clean.

**Fig. 2.**
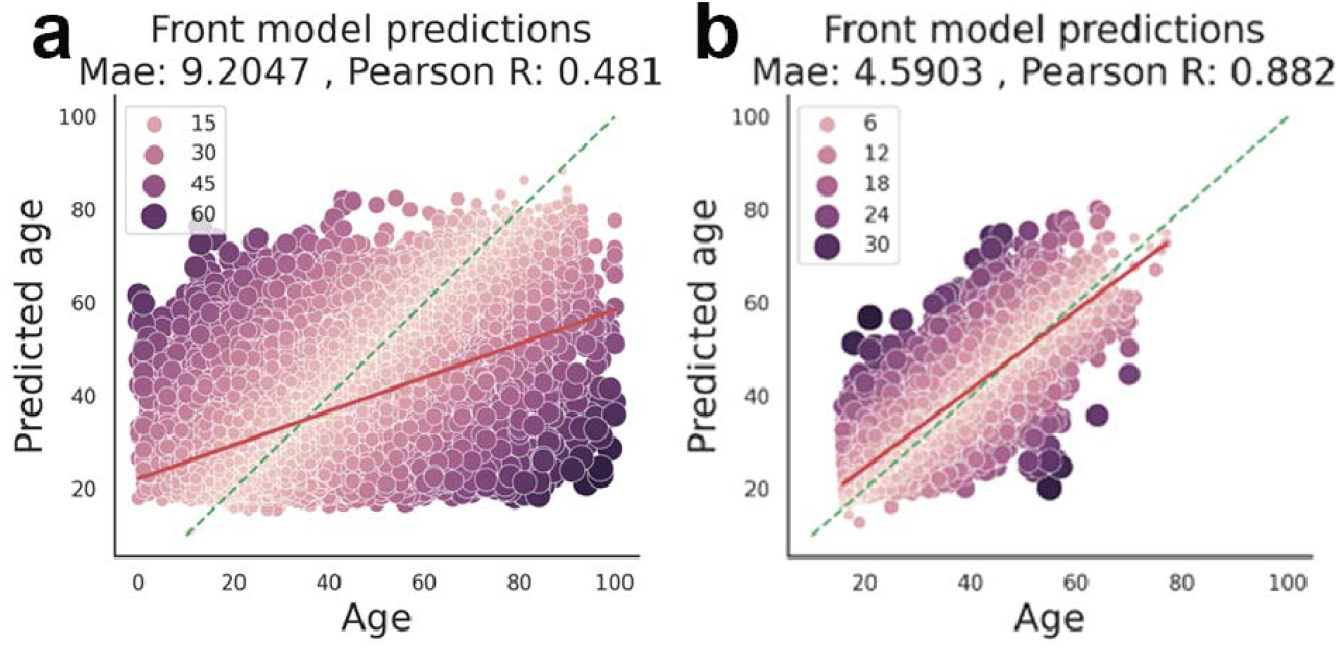
**a**, Age prediction of the *Front model* on the IMDB-Clean dataset. **b**, Age prediction of the *Front model* on the MORPH-2 dataset.

Overall, we conclude that accurate age prediction is possible using side photos despite the smaller facial area compared to front-facing photos. The age prediction performance further improved by using two images from one person at the same time, captured from different angles, front and side.

### The age acceleration based on front-view face photos correlated with the age acceleration based on side-view face photos

We examined the correlations between the different models applied on the test set of the Prisoner dataset (Fig. 3). Age prediction of the different models correlated well with each other, not surprisingly, as all of them correlated with age (r > 0.91). Age acceleration, which is the deviation of the predicted age from the trend, did not correlate with age, allowing the comparability of different age groups in age acceleration analyses. However, the age acceleration of the *Front model* significantly correlated with the age acceleration of the *Side model* (r = 0.49), meaning that subjects that were aging faster based on their frontal face photo generally were aging faster based on their side-view photo too (Fig. 3). The age acceleration of the *Front model* significantly correlated with the age acceleration of the *Side model* (r = 0.28) in the external test set (the SCFace dataset) too (Fig. 4).

**Fig. 3.**
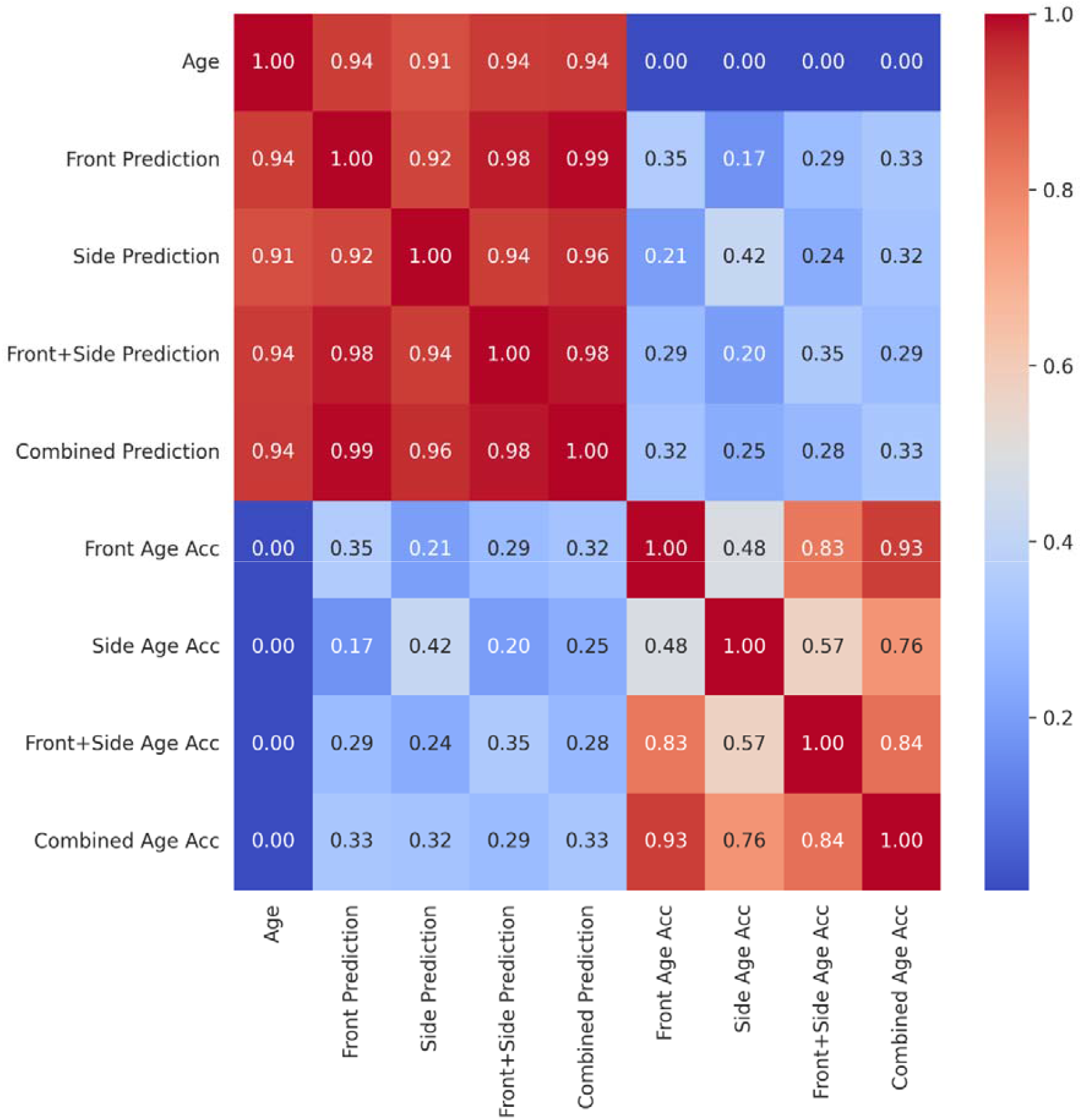
All vs all correlation analysis of age, predicted age, and age acceleration of the four face photo-based aging clock models on the testing set of the Prisoner dataset.

**Fig. 4.**
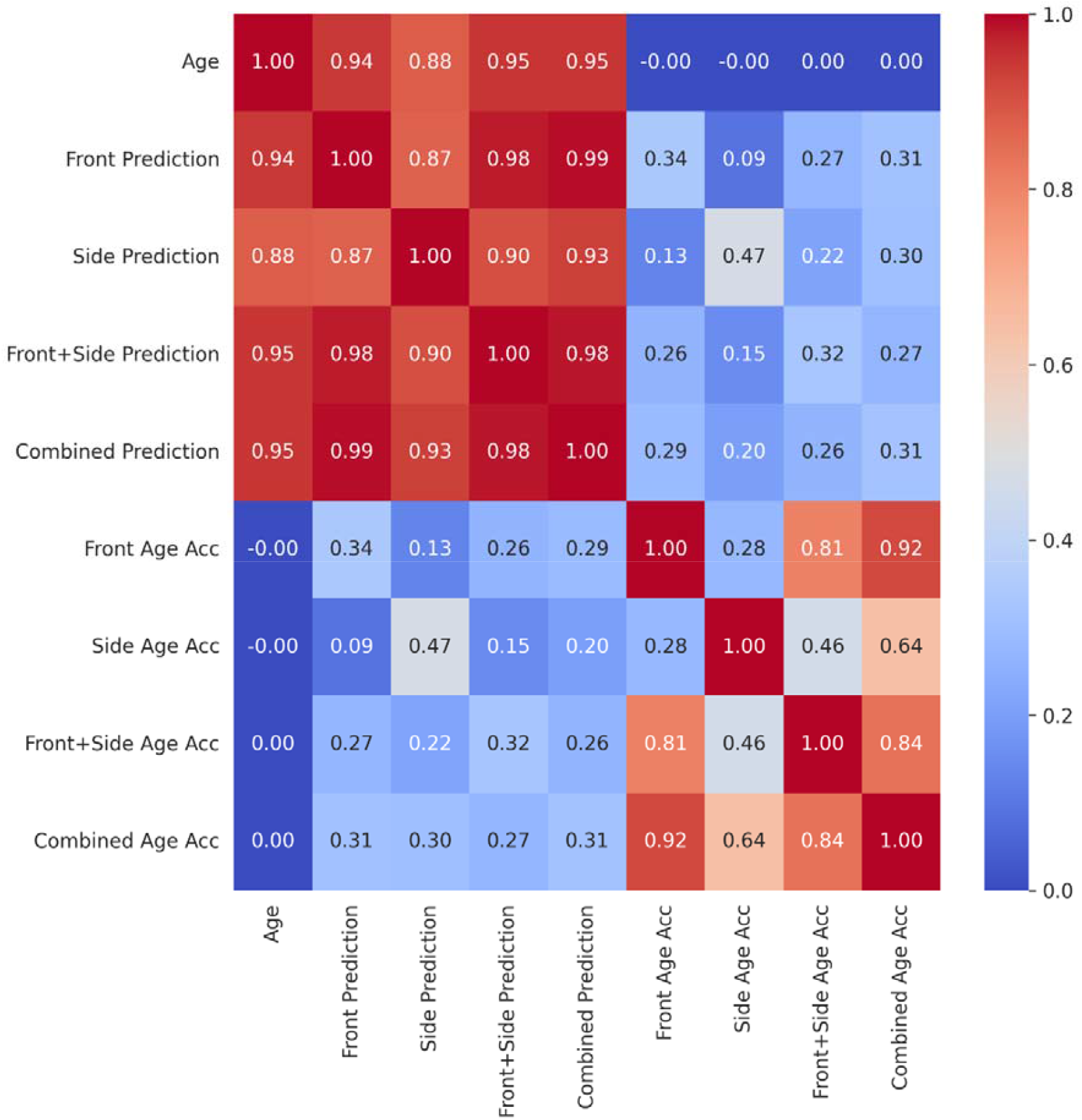
All vs all correlation analysis of age, predicted age, and age acceleration of the four face photo-based aging clock models on the SCface dataset.

We also observed that the correlation of age accelerations between the *Front* and *Front + Side model* was higher than the correlation of age accelerations between the *Side* and *Front + Side* model. The same was true for the *Combined model*. We concluded that while age prediction using side-view face photos showed comparable accuracy as front-view face photos, the frontal-view photo became more important for the two-image models.

### The *Side model* handles the rotation of the face better compared to the *Front model*

The SCface dataset contains nine pictures from different angles of every individual from left to right side in equal steps by 22.5 degrees. The different angles are labeled as “frontal”, “L1” to “L4”, and “R1” to “R4”, as the angle increases from the frontal picture (Fig. 5a). When we tested the *Front model* on the SCface dataset in all available angles, we observed that the age prediction became less accurate as the angle increased starting from the front-facing image (Fig. 5bc, Supplementary Table S2). Interestingly, the *Side model* handled the rotation of the face better, providing more consistent results (the MAE was between 2.47 and 10.48 for the *Front model* while between 2.35 and 3.87 for the *Side model*). Furthermore, rotating the face provided more accurate age prediction at almost every angle, except for the front-facing and slightly rotated (R1, L1) images. As the angle decreased, the MAE of the *Side model* increased only to a small extent, much less compared to the *Front model*.

**Fig. 5.**
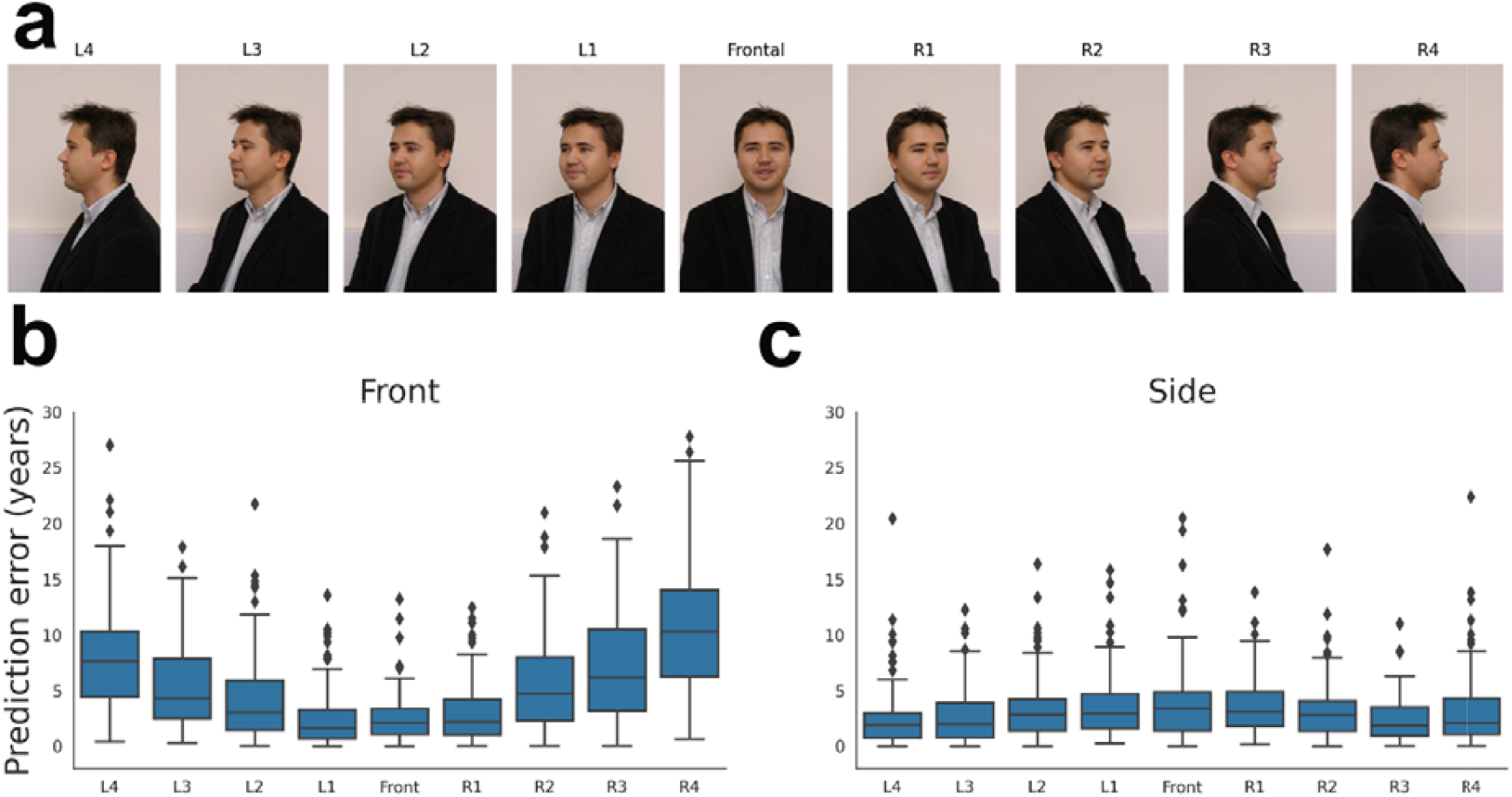
**a**, Example images from the SCface dataset (adapted from the website of the SCface database https://www.scface.org/)^19^. **b**, Absolute error of the *Front model* applied on different face angles on the SCface dataset. **c**, Absolute error of the *Side model* applied on different face angles on the SCface dataset.

Then we tested the *Side model* exclusively on the front images of the Prisoner-, IMDB Clean-, and MORPH-2 datasets (Supplementary Table S3) and observed slightly worse results compared to the *Front model*. In contrast, when we tested the *Front model* exclusively on side-view photos, the *Front model* performed much worse than the *Side model* (Supplementary Table S4). In summary, the *Side model* performed much better on the frontal-view face photos than the *Front model* on the side-view face photos. An explanation of this observation can be that the *Side model* may focus on features that are completely covered in all-angle photos, while the *Front model* may focus on the nose-mouth-eyes areas that are not completely covered in the side photos.

## Discussion

Here, we developed deep learning age prediction models using side-view face photos. We conclude that it is possible to predict age from side-view face photos with almost the same accuracy as using front-view face photos. Our analysis showed that a benefit of the side-view approach is that it handles face rotation better than the traditional frontal models.

We also showed that two photos at different angles (front and side) improved age prediction and may provide a better approach for determining biological age. So far, only a few studies reported age predictions based on multimodal data (i.e, multiple types of measurements of the same individual at the same time). Wang et al. trained age prediction models using 3D facial, tongue, and retinal images from healthy subjects and demonstrated that using a fusion of the three image modalities achieved the most accurate predictions^16^. Recently, we developed aging clocks using the methylation and gene expression features at the same time, and found that the most accurate models tended to use exclusively methylation features^20^. Future studies needs to explore the benefit of the different data modelities in the measuring of biological age.

In summary, we showed that two photos at different angles can improve age prediction and may provide a better approach to determining biological age for personalized medicine and rejuvenation studies.

## Methods

### The Prisoner dataset

We used the Kaggle IDOC (Illinois Department of Corrections) dataset (https://www.kaggle.com/datasets/davidjfisher/illinois-doc-labeled-faces-dataset) and tabular data with rich annotations from the Illinois Prison Population Data Sets (https://idoc.illinois.gov/reportsandstatistics/populationdatasets.html). The dataset contained mugshots (i.e., photographic portraits of each subject from the shoulders up with plain background, one front-view photo, and one side-view photo) of individuals who were in prison at the end of 2018. The mugshots are taken at the date of admission to the prison but the photos are updated periodically. To decrease the chance that the mugshots were updated since the date of admission, we used only the mugshots of individuals with an admission date between 2015 and 2018. We used additional mugshots directly downloaded from the website of the Illinois Prison Population Data Sets (https://idoc.illinois.gov/reportsandstatistics/populationdatasets.html) from individuals who were in prison in July of 2024, and their admission dates were between 2021 and 2024. Metadata was also available through the Illinois Prison Population Data Set website. We assigned ages to the photos using the formula Age = Admission date - Birth date. In summary, we collected high-quality mugshots of 51,149 individuals, 47,642 of males, 3,507 of females. The distribution of ethnicity was 63 of Amer Indian, 156 of Asian, 116 of Bi-racial, 27,617 of Black, 5,638 of Hispanic, 17,500 of White, and 59 of Unknown ethnicity. The youngest age was 17 years, and the oldest age was 83 years.

### Training and testing of the age prediction models

We split the data into training, validation, and testing sets by using 0.6, 0.2, and 0.2 ratio, respectively, and the removed those pictures when multiple pictures were taken of one person. After this, the training set contained 32,577 images, the validation set contained 9,281 images, and the test set contained 9,291 images. All of the images were cropped using the Retinaface face detector^21^ followed by padding to a square and resized to 224×244 pixels. Then we trained a Res-Net 50 model on the training dataset by modifying the final layer to a single-layer neuron. We trained separate models using (i) only front face photos (*Front model*), (ii) only side view face photos (*Side model)*, and (iii) both front and side view face photos (*Front + Side model)*. We constructed another model by combining the front and the side models by using an Ordinary Least Squares method on the validation set, and tried to predict the age from the Front and Side model predictions. We named the output of this model as *Combined Model* with the formula (predicted age) = −0.7285+ 0.7467 □ (Front model predictions) + 0.2951 □ (Side model predictions), and after this we tested this model on the Test Set and the Sc Face dataset.

### External testing datasets

We used the SCface dataset (https://www.scface.org/) for external testing^22^, containing high-quality mugshots of 130 people (116 males and 14 females) captured from 9 different angles. The database contained additional annotations about gender, beard, moustache, and glasses.

We also used another independent mugshot dataset, the MORPH-2 (https://www.kaggle.com/datasets/chiragsaipanuganti/morph)^23^, for an additional external testing of our *Front model*. The MORPH-2 dataset contained 55,134 images of subsets aged between 16 and 77 years.

We tested our *Front model* on the IMDB-Clean dataset (https://github.com/yiminglin-ai/imdb-clean?tab=readme-ov-file), which was an “in-the-wild” dataset, meaning that the photos were taken in real-life scenarios^24^. This dataset consisted of 221,055 images.

## Statistics

To measure the performance of our models, we used R^2^ score, Pearson R, Mean average error (MAE), and Cumulative Score 3 CS_3_(%) (i.e., the percentage of the predictions had less than 3 years of absolute error). We used p-values to determine whether the correlation is statistically significant. Significance levels are marked using asterisks, ns : p > 0.05, * : 0.01 < p ≤ 0.05, ** : 0.001 < p ≤ 0.01, *** : p ≤ 0.001, ****: p ≤ 0.0001.

## Data and code availability

The Python code for testing the models is available on GitHub: https://github.com/bdbotond/dual_face_age_pred.

## Author contributions

BB-D had the initial idea for using side-view photos for age prediction. BB-D found useful datasets for the study, developed the models and conducted the data analysis. CK supervised the project. Both authors wrote the manuscript.

### Acknowledgments

The project was supported by the European Union project RRF-2.3.1-21-2022-00004 within the framework of the Artificial Intelligence National Laboratory. The project was also supported by the National Research, Development and Innovation Office – NKFIH, FK-146113.

## SUPPLEMENTARY TABLES

**Supplementary Table S1.**

| Dataset | $R^2$ Score | Pearson R | MAE | $CS_3(\%)$ |
| --- | --- | --- | --- | --- |
| Results of the model that uses only the front images |  |  |  |  |
| Prisoner front images | 0.87 | 0.94 | 3.11 | 0.8 |
| SCface front images | 0.87 | 0.94 | 2.48 | 0.93 |
| MORPH-2 | 0.72 | 0.88 | 4.59 | 0.62 |
| IMDB-Clean | 0.14 | 0.48 | 9.23 | 0.37 |
| Results of the model that uses only the side images |  |  |  |  |
| Prisoner side images | 0.82 | 0.91 | 3.69 | 0.73 |
| SCface side images | 0.76 | 0.88 | 3.05 | 0.82 |
| Results of the model that uses the front images and the side images |  |  |  |  |
| Prisoner two images | 0.88 | 0.93 | 3.01 | 0.82 |
| SCface two images | 0.89 | 0.95 | 2.49 | 0.89 |
| Results of the regression model that uses a regression with the front and the side model |  |  |  |  |
| Prisoner two images | 0.89 | 0.94 | 2.88 | 0.83 |
| SCface two images | 0.89 | 0.94 | 2.31 | 0.91 |

**Supplementary Table S2.**

| MAE of the Front model and the Side model from different angles |  |  |
| --- | --- | --- |
| Angle | MAE (Front model) | MAE (Side model) |
| L1 | 2.47 | 3.56 |
| L2 | 4.18 | 3.3 |
| L3 | 5.27 | 2.62 |
| L4 | 8.18 | 2.47 |
| R1 | 2.98 | 3.5 |
| R2 | 5.74 | 3.14 |
| R3 | 7.02 | 2.35 |
| R4 | 10.48 | 3.05 |
| frontal | 2.48 | 3.87 |

**Supplementary Table S3.**

| Side model results compared to the Front model results on frontal images |  |  |
| --- | --- | --- |
|  | Side model | Front model |
| Dataset | MAE | MAE |
| Prisoner front images | 6 | 3.14 |
| SCface front images | 3.87 | 2.48 |
| MORPH-2 | 7.21 | 4.59 |
| IMDB-Clean | 10.17 | 9.23 |

**Supplementary Table S4.**

| Front model results compared to the Side model results on side images |  |  |
| --- | --- | --- |
|  | Front model results on the side images | Side model results on the Side images |
| Dataset | MAE | MAE |
| Prisoner front images | 8.93 | 3.72 |
| SCface front images | 10.48 | 3.05 |

